# *MoDentify*: a tool for phenotype-driven module identification in multilevel metabolomics networks

**DOI:** 10.1101/275057

**Authors:** Kieu Trinh Do, David J.N.-P. Rasp, Gabi Kastenmüller, Karsten Suhre, Jan Krumsiek

## Abstract

**Summary:** Metabolomics is an established tool to gain insights into (patho)physiological outcomes. Associations of metabolism with such outcomes are expected to span functional modules, which are defined as sets of correlating metabolites that are coordinately regulated. Moreover, these associations occur at different scales, from entire pathways to only a few metabolites, which is an aspect that has not been addressed by previous methods. Here we present *MoDentify,* a freely available R package to identify regulated modules in metabolomics networks at different layers of resolution. Importantly, *MoDentify* shows higher statistical power than classical association analysis. Moreover, the package offers direct visualization of results as interactive networks in Cytoscape. We present an application example using a complex, multifluid metabolomics dataset. Owing to its generic character, the method is widely applicable to any dataset with a phenotype variable, a data matrix, and optional pathway annotations.

**Availability and Implementation:** *MoDentify* is freely available from GitHub: https://github.com/krumsiek/MoDentify

The package vignette contains a detailed tutorial of the analysis workflow.

## 1 Introduction

Associations with phenotypic parameters and clinical endpoints in large-scale, heterogeneous metabolomics datasets are complex. They typically span entire functional modules, which are defined as groups of correlating molecules that are functionally coordinated, coregulated, or generally driven by a common biological process (Mitra *et al,* 2013).

The systematic identification of modules is often based on networks, where nodes correspond to the molecules under investigation, and edges represent the correlations or associations between two molecules. Modules are commonly identified as highly connected parts of the network that contain nodes that are coordinately associated with a given phenotype.

Systematic module identification algorithms are well established for various types of *omics* data (Polanski *et al,* 2014; Chuang *et al,* 2007; May *et al,* 2016; Martignetti *et al,* 2016). However, they have scarcely been applied to metabolomics data. Moreover, none of these methods consider that phenotype associations can occur at different scales, ranging from global associations spanning entire pathways or even sets of pathways (e.g., “dense” associations between metabolomics and gender or BMI), to localized associations with only a few metabolites (e.g., “sparse” associations between metabolomics and insulin-like growth-factor I levels or asthma) (Do *et al,* 2017). For sparse associations, the identification and interpretation of modules is usually straightforward. However, modules for dense phenotype associations at the metabolite level are challenging to interpret due to their overwhelming number. To facilitate interpretation, the plethora of information at the fine-grained metabolite level can be condensed to a hierarchically superordinate level, such as a pathway network. Here, nodes correspond to entire pathways, edges represent pathway relationships, and modules reflect phenotype-associated processes covering sets of pathways.

We have recently introduced a module identification algorithm for multifluid metabolomics data (Do *et al*, 2017). The approach was applied to blood concentrations of insulin-like growth factor (IGF-I) and gender as examples of sparse and dense phenotype associations, respectively. We here present *MoDentify*, a free R package implementing the approach for general use. In particular, *MoDentify* offers (i) the estimation of data-driven networks based on Gaussian graphical models (GGMs), (ii) module identification at both fine-grained metabolite level and more global pathway levels, and (iii) visualization of the identified modules in an interactive network through Cytoscape (Shannon *et al*, 2003). *MoDentify* increases statistical power compared with classical association analysis due to the reduction of statistical noise and can easily be applied to any type of quantitative data because of its generic character.

## 2 Description

*MoDentify* identifies network-based modules that are highly affected by a phenotype of interest. The underlying network is either directly inferred from the data at the single metabolite or pathway level (see below) or can be provided from an external source.

### Network inference

*MoDentify* estimates either classical Pearson correlation networks or GGMs using the *GeneNet* R package (Opgen- Rhein & Strimmer, 2007). GGMs are based on partial correlations, which represent associations between two variables corrected for all remaining variables in multivariate Gaussian distributions (Krumsiek *et al,* 2011). An important property of GGMs compared with Pearson correlation networks is their sparsity, because only direct correlations are included. At the fine-grained level, the GGM consists of nodes corresponding to metabolites and edges representing significant partial correlations between two nodes after multiple testing correction. At the pathway level, the GGM consists of nodes corresponding to entire pathways (sets of metabolites), whereas edges represent significant partial correlations between two pathways. To estimate a correlation network between pathways, representative values are computed for each pathway (see **Pathway representation**). Alternatively, a network from an external source can be provided. Importantly, all nodes in the network must be measured in the given dataset.

### Pathway representation

As stated above, in addition to regular network inference, *MoDentify* can build a network of interacting pathways. To this end, a new variable is defined as a representative for each pathway, which aggregates the total abundance of metabolites from the pathway into a single value. *MoDentify* provides two approaches for pathway representation:

1. *eigenmetabolite* approach: For each pathway, a principal component analysis (PCA) is performed after scaling all metabolites to a mean of 0 and a variance of 1. The first principle component - also termed *eigenmetabolite -* is used as a representative value for the entire set of variables in the pathway (Langfelder and Horvath 2007).
2. *average* approach: All variables are first scaled to mean 0 and variance 1. Subsequently, the pathway representative is calculated as the average of all variable values in the pathway.

*MoDentify* computes the amount of explained variances explained by each eigenmetabolite per pathway to facilitate the choice between these two approaches. If explained variances are high, the *eigenmetabolite* approach should be used; otherwise, the *average* approach might be the more appropriate choice.

### Module identification

To identify functional modules, *MoDentify* uses a score maximization approach. Given a network, a scoring function, and a starting node (seed node) as the initial candidate module, the algorithm identifies an optimal module by score maximization. For a given candidate module, the following procedure is performed: Each neighboring node of the module is subsequently added to the candidate module, and the score of the extended module is calculated (see Module scoring). The neighbor resulting in the highest score improvement is finally added to the candidate module, if the new module score is higher than the score of each of its single components. The procedure is repeated until no further score improvements can be made, yielding the optimal module for the given seed node. In an optional consolidation step, overlapping optimal modules from different seed nodes are combined into one module, which is reevaluated by the scoring function.

### Module scoring

The score of a candidate module is obtained from the multivariable linear regression model

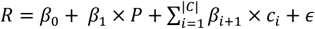

where *R* is the module representative defined by the *eigenmetabolite* or *average* approach (see **Pathway representation**), *β*_*0*_ is the intercept, *β*_*i*_ are the regression coefficients for the respective independent variables, *P* is the phenotype of interest, *C* is an optional set of covariates *c*_*t*_, and *e* is a normally distributed error term. The module score is then defined as the negative log-transformed p-value of *β*_*1*_. The significance of the modules is assessed by correcting for the total number of nodes in the underlying network.

### Module visualization

*MoDentify* offers visualization of the identified modules within an interactive network in the open source software Cytoscape (Shannon *et al*, 2003). The network contains different node colors and node sizes, which depict the membership of a node in a certain module and its association with the given phenotype from a classical, single-molecule association analysis, respectively. Moreover, significance of the phenotype association is indicated by diamond-shaped nodes. In addition to returning R data structures and producing flat-file results, one of the main advantages of our module visualization is the direct call of Cytoscape from within R *via* the *RCytoscape* package (Shannon *et al,* 2013) for external visualization, without cumbersome exporting of data files from R and re-importing them into Cytoscape.

## 3 Application example

We demonstrate the easy use of *MoDentify* on plasma, urine, and saliva metabolomics data from the Qatar Metabolomics Study on Diabetes (QMDiab) (Mook-Kanamori *et al,* 2014), aiming to identify functional modules associated with type 2 diabetes (T2D). The multifluid dataset comprises mass spectrometry-based metabolomics measurements for 190 diabetes patients and 184 healthy controls of Arab and Asian ethnicities aged 17-81 years. The dataset consists of 1524 metabolites. For each metabolite, two levels of pathway annotation are available. The preprocessed QMDiab data (normalized, log-transformed, missing values handled, and scaled) are integrated within the *MoDentify* package. The dataset is also available from the following figshare repository via the following link https://doi.org/10.6084/m9.figshare.5904022.

*MoDentify* was applied to the QMDiab dataset at both metabolite and pathway levels. The following code with default parameters produces a list of metabolite modules associated with T2D, as well as interactive visualization of the modules in the underlying network in Cytoscape (Figure 1A). Here, we only show code for the application of *MoDentify* at the metabolite level. Code for application at the pathway level (Figure 1B) can be found in the package vignette, available from the GitHub repository.

**Figure 1.**
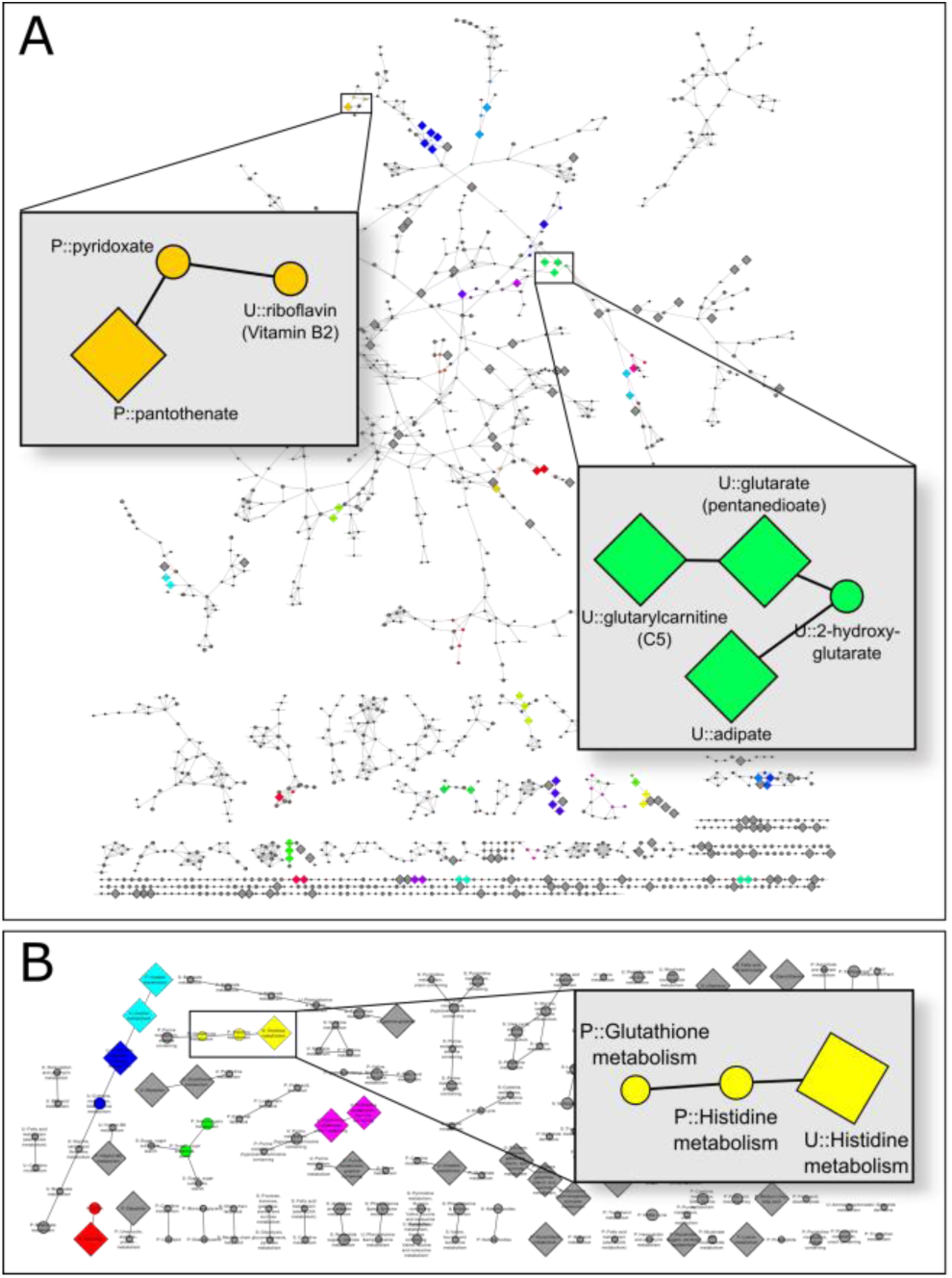
Visualization of identified modules for type 2 diabetes. The metabolomics networks with embedded modules at metabolite level (A) and pathway level (B) are screenshots of the interactive versions in Cytoscape produced by *MoDentify.* Zoom-ins have been added to highlight examples for *MoDentify’s* increased statistical power and its ability to extract biologically valuable insights. Round nodes correspond to metabolic entities not significantly associated with T2D when considered alone. Diamond nodes represent metabolic entities significantly related to T2D.

~~~
# *Load MoDentify* library(MoDentify)
# *Network inference*
met.graph <- generate.network(data = qmdiab.data, annotations = qmdiab.annos)
# *ModuLe identification*
modules.summary <- identify.modules(graph = met.graph, data = qmdiab.data, annotations = qmdiab.annos, phenotype = qmdiab.phenos$T2D)
# *ModuLe visualization*
draw.modules(graph = met.graph, summary = modules.summary)
~~~

By default, `generate.network` estimates partial correlations between metabolites and assigns edges using a significance threshold of *α* = 0.05 after Bonferroni multiple testing correction. `identify.modules` searches network modules for the given phenotype, where the default module representation approach is the *average* approach, *α* = 0.05 is set for significance filtering, and Bonferroni multiple testing correction is applied. The output structure modules. summary contains a list of modules with their components and scores, which can be visualized within an interactive network in Cytoscape using `draw.modules`.

*MoDentify* identified 36 modules for T2D at the metabolite level (Figure 1A). Many of these modules consist of metabolites that are not significantly associated with T2D if considered alone. However, in interplay with other metabolites, they form a module that is more strongly associated with T2D than all of its single components. This increased statistical power in *MoDentify* can be attributed to the reduction of statistical noise when aggregating module components and allows the detection of links between metabolites and phenotype that would have been missed with classical association analysis. *MoDentify* found several modules containing metabolites from at least two fluids. For instance, one module (orange in Figure 1A) comprises the three vitamin B derivatives plasma pantothenate (vitamin B_5_) and pyridoxate (vitamin B_6_), and urine riboflavin (vitamin B_2_). Although pyridoxate and riboflavin are not related to T2D when analyzed alone, they form a module in combination with pantothenate that is significantly associated with the phenotype. This module corroborates previous observations that vitamin B levels in blood and urine are associated with T2D (Nix *et al,* 2015; Unoki-Kubota *et al,* 2010; Valdes-Ramos *et al,* 2015). In addition, the results indicate that not only the concentration levels in blood and urine but also exchange processes between the two fluids are linked to T2D as well. At the pathway level (Figure 1B), six modules were detected. These modules show the interplay of multiple pathways in diabetes. For instance, one module comprises plasma metabolites from glutathione and histidine metabolism and urinary metabolites from histidine metabolism (yellow in Figure 1B). Although histidine and glutathione were shown to be related to diabetes in previous studies (Kimura *et al,* 2013; Sekhar *et al,* 2011), the identified module suggests that histidine and glutathione metabolism as well as the secretion of histidine derivatives might be part of the same process in T2D.

## 4 Conclusion

To the best of our knowledge, *MoDentify* implements the first approach for the systematic identification of phenotype-driven modules at different layers of resolution. To this end, the algorithm allows the estimation of data-driven networks based on Pearson or partial correlations. Optionally, a network from an external source can be provided. To facilitate result interpretation for different scales of phenotype associations, *MoDentify* enables the module search at both fine-grained metabolite level and more global pathway levels. Owing to the increased statistical power of the approach, novel links between clinical parameters and molecular levels can be detected. We presented an applicationexample using a complex multifluid metabolomics dataset, but owing to its generic character, this approach can be applied for any quantitative dataset.

## Acknowledgments

We wish to thank the study participants and research team of the QMDiab study for their contributions. The QMDiab study was approved by the Institutional Review Boards of HMC and WCM-Q under research protocol number 11131/11. All study participants provided written informed consent.

## Funding

KD is supported by a grant (Grant no. 01ZX1313C, project e:Athero-MED) from the German Federal Ministry of Education and Research (BMBF). JK has received funding from the European Union’s Seventh Framework Program (FP7-Health-F5- 2012) under grant agreement no. 305280 (MIMOmics). GK is supported by funding from the National Institute of Aging (NIH/NIA) under grant no. 1RF1AG057452-01 (Metabolic Network Analysis of Biochemical Trajectories in Alzheimer’s Disease) and from Qatar National Research Fund grant NPRP8-061-3-011 (Rational Metabolic Engineering as a New Tool for Targeted Intervention in Human Metabolism). All statements are made on behalf of the authors and do not necessarily represent the views of the research agencies. KS is supported by Biomedical Research Program funds at Weill Cornell Medical College in Qatar, a program funded by the Qatar Foundation.

*Conflict of Interest:* none declared.

## References

Chuang, H.Y. et al. (2007) Network-based classification of breast cancer metastasis. Mol. Syst. Biol., 3, 140

Do, K.T. et al. (2017) Phenotype-driven identification of modules in a hierarchical map of multifluid metabolic correlations. NPJ Syst. Biol. Appl., 3, 28

Kanehisa, M. et al.(2012) KEGG for integration and interpretation of large-scale molecular data sets. Nucleic Acids Res., 40, D109–114

Kimura, K. et al. (2013) Histidine Augments the Suppression of Hepatic Glucose Production by Central Insulin Action. Diabetes 62: 2266–2277

Krumsiek, J. et al. (2011) Gaussian graphical modeling reconstructs pathway reactions from high-throughput metabolomics data. BMC Syst. Biol., 5, 21

Martignett, L. et al. (2016) ROMA: Representation and quantification of module activity from target expression data. Front. Genet., 7, 18

May, A. et al. (2016) Metamodules identifies key functional subnetworks in microbiome-related disease. Bioinforma. Oxf. Engl., 32, 1678–1685

Mitra K, et al. (2013) Integrative approaches for finding modular structure in biological networks. Nat. Rev. Genet., 14, 719–732

Mook-Kanamori DO, et al. (2014) 1,5-anhydroglucitol in saliva is a noninvasive marker of short-term glycemic control. J. Clin. Endocrinol. Metab., 99, E479–E483

Nix WA, et al. (2015) Vitamin B status in patients with type 2 diabetes mellitus with and without incipient nephropathy. Diabetes Res. Clin. Pract., 107, 157165

Opgen-Rhein, R. and Strimmer, K. (2007) From correlation to causation networks: a simple approximate learning algorithm and its application to high-dimensional plant gene expression data. BMC Syst. Biol., 1, 37

Polanski K, et al. (2014) Wigwams: identifying gene modules co-regulated across multiple biological conditions. Bioinformatics 30, 962–970

Sekhar RV, et al. (2011) Glutathione synthesis is diminished in patients with uncontrolled diabetes and restored by dietary supplementation with cysteine and glycine. Diabetes Care 34, 162–167

Shannon, P. et al. (2003) Cytoscape: a software environment for integrated models of biomolecular interaction networks. Genome Res. 13: 2498–2504

Shannon, P.T. et al. (2013) RCytoscape: tools for exploratory network analysis. BMC Bioinformatics 14, 217

Swainston, N. et al. (2016) Recon 2.2: from reconstruction to model of human metabolism. Metabolomics 12, Available at: http://www.ncbi.nlm.nih.gov/pmc/articles/PMC4896983/ [Accessed August 7, 2017]

Unoki-Kubota, H. et al. (2010) Pyridoxamine, an inhibitor of advanced glycation end product (AGE) formation ameliorates insulin resistance in obese, type 2 diabetic mice. Protein Pept. Lett., 17. p1177–1181

Valdés-Ramos, R. et al. (2015) Vitamins and type 2 diabetes mellitus. Endocr. Metab. Immune Disord. Drug Targets 15, p54–63

